# Into the Evening: Complex Interactions in the *Arabidopsis* circadian clock

**DOI:** 10.1101/068460

**Authors:** He Huang, Dmitri A. Nusinow

## Abstract

In *Arabidopsis thaliana*, an assembly of proteins named the evening complex (EC) has been established as an essential component of the circadian clock with conserved functions in regulating plant growth and development. Recent studies identifying EC-regulated genes and EC-interacting proteins have expanded our understanding of EC function. In this review, we focus on new progress uncovering how the EC contributes to the circadian network through the integration of environmental inputs and the direct regulation of key clock genes. We also summarize new findings of how the EC directly regulates clock outputs, such as day-length dependent and thermoresponsive growth, and provide new perspectives on future experiments to address unsolved questions related to the EC.

## The evening complex (EC): a key component of the plant circadian clock network

The circadian clock is an endogenous time-keeping mechanism with a period of approximately 24 hours that is found in all domains of life [1](Box 1). Circadian oscillators function to coordinate internal physiology with external environmental cues, such as sunrise and sunset, to provide an adaptive advantage [2-5]. Circadian clocks can be observed in organisms as persistent 24-hour rhythms in constant environments, as was described in 1729 by de Mairan who observed that daily sleep movements in mimosa leaves continued in constant darkness [6]. In eukaryotes, clock networks are composed of multiple interlocked transcriptional-translational feedback loops (TTFLs) [7]. While the genes constituting TTFLs are largely distinct between plants, animals, and fungi, the common architecture and mechanisms of the circadian systems ensure that insights can shared between diverse organisms.

#### Box 1 Basic Concepts of Circadian Clock Systems

Circadian clock systems are found in all domains of life [1] and share three common modules: inputs, circadian oscillators and outputs. Input pathways convert external cues (also known as zeitgebers, German for time givers), such as ambient light and temperature, to circadian oscillators to reset and synchronize the clock with the local environment (entrainment). In eukaryotes, circadian oscillators are cell-autonomous pacemakers with a period of ˜24 hours composed of mostly transcription factors. Circadian clocks generate and sustain rhythmicity for long periods even in the absence of environmental cues (i.e. free-running conditions, such as constant light and temperature conditions). In addition, circadian clocks also maintain a 24 hour period regardless of changes in ambient temperature, a phenomena known as temperature compensation. Upon receiving timing cues from input pathways, oscillators link to various output processes to rhythmically regulate the levels of genes, proteins and metabolites, allowing organisms to anticipate and adapt to the changing environments, such as seasonal changes in day length (photoperiod) and temperature. Output pathways under circadian clock regulation exhibit a rhythmic response to a constant input, a phenomenon defined as “gating”. Gating by the clock allows for outputs to occur at specific times of day or to coordinate with environmental changes such as day/night transitions.

In the model plant *Arabidopsis thaliana*, the use of genetic screens and non-invasive luciferase reporters has led to the identification of dozens of core clock or clock-associated components. These genes form a complex network of morning-, afternoon- and evening-phased transcription and translation feedback loops that generate 24-hour oscillations (Box 2 and Figure 1, **Key Figure**) [8-11]. Within this network, a tripartite protein complex called the evening complex (EC) is an essential component of the evening transcription loop [8, 9, 12]. The naming of the EC is derived from the observation that both gene expression and protein levels of all three EC components, namely EARLY FLOWERING 3 (ELF3), EARLY FLOWERING 4 (ELF4) and LUX ARRHYTHMO (LUX, also known as PHYTOCLOCK1), peak at dusk [12-17]. In contrast to other clock genes, loss-of-function mutation in any of the EC components (*elf3, elf4* or *lux*) causes an arrhythmic phenotype [13, 16-18]. This clock arrhythmia is accompanied by many phenotypes, such as inappropriate cellular elongation in response to environmental cues and early flowering regardless of day length [12-17, 19-22]. These results collectively demonstrate the indispensable role of the EC in maintaining circadian rhythms and coordinating growth and development.

#### Box 2 Introduction of the TTFLs in *Arabidopsis thaliana*

Knowledge of the plant circadian clock mostly comes from studying the model *Arabidopsis thaliana*. Accumulated studies using genetic, molecular and biochemical approaches have identified many components and enhanced our understanding of the oscillatory mechanism in plants [9]. In *A. thaliana*, the circadian clock is currently described as containing multiple inter-connected transcriptional-translational feedback loops (TTFLs) [8](Figure 1).

The morning loop is composed of two MYB-domain-containing transcription factors CIRCADIAN CLOCK ASSOCIATED 1 (CCA1) and LATE ELONGATED HYPOCOTYL (LHY), which form a heterodimer and function synergistically to regulate the expression of other clock genes [101]. CCA1 and LHY positively regulate the expression of the morning-phased *PSEUDO-RESPONSE REGULATOR 9 (PRR9)* and afternoon-phased PRR7 [102, 103], while negatively regulating another two afternoon-phased genes *PRR5* and *GIGANTEA (GI)* [33, 34], and several evening-phased genes, such as *EARLY FLOWERING 3 (ELF3), EARLY FLOWERING4 (ELF4), LUX ARRHYTHMO (LUX)* and *TIMING OF CAB EXPRESSION1* (TOC1, also known as PRR1) [17, 29, 33, 104]. PRR9, PRR7, and PRR5 feedback and repress the expression of *LHY* and *CCA1* during the day, allowing the induction of evening genes [99, 103]. In addition, another morning-phased REVEILLE8 (RVE8) induces evening-expressed clock genes *TOC1, PRR5, ELF4,* and *LUX* in the afternoon (˜8 hours after dawn) [37]. In the early evening, TOC1 then negatively regulates CCA1 and LHY through direct association with their promoters [32, 105]. A multi-protein complex comprised of *ELF3, ELF4* and *LUX,* named the evening complex (EC), represses *PRR9* in the late evening to allow CCA1 and LHY levels to rise before dawn [12, 23-25, 41].

It has been found that the circadian clock regulates various output pathways in *A. thaliana*, including photosynthesis, growth, disease resistance, starch metabolism and phytohormone pathways [47, 106, 107]. Comparative genomics analysis has found that circadian clock components are broadly retained after genome duplication events, reflective of positive selection [108]. Thus, the circadian clock system is a target for manipulation that could potentially lead to improvement of plant fitness and yield of crops.

**Figure 1.**
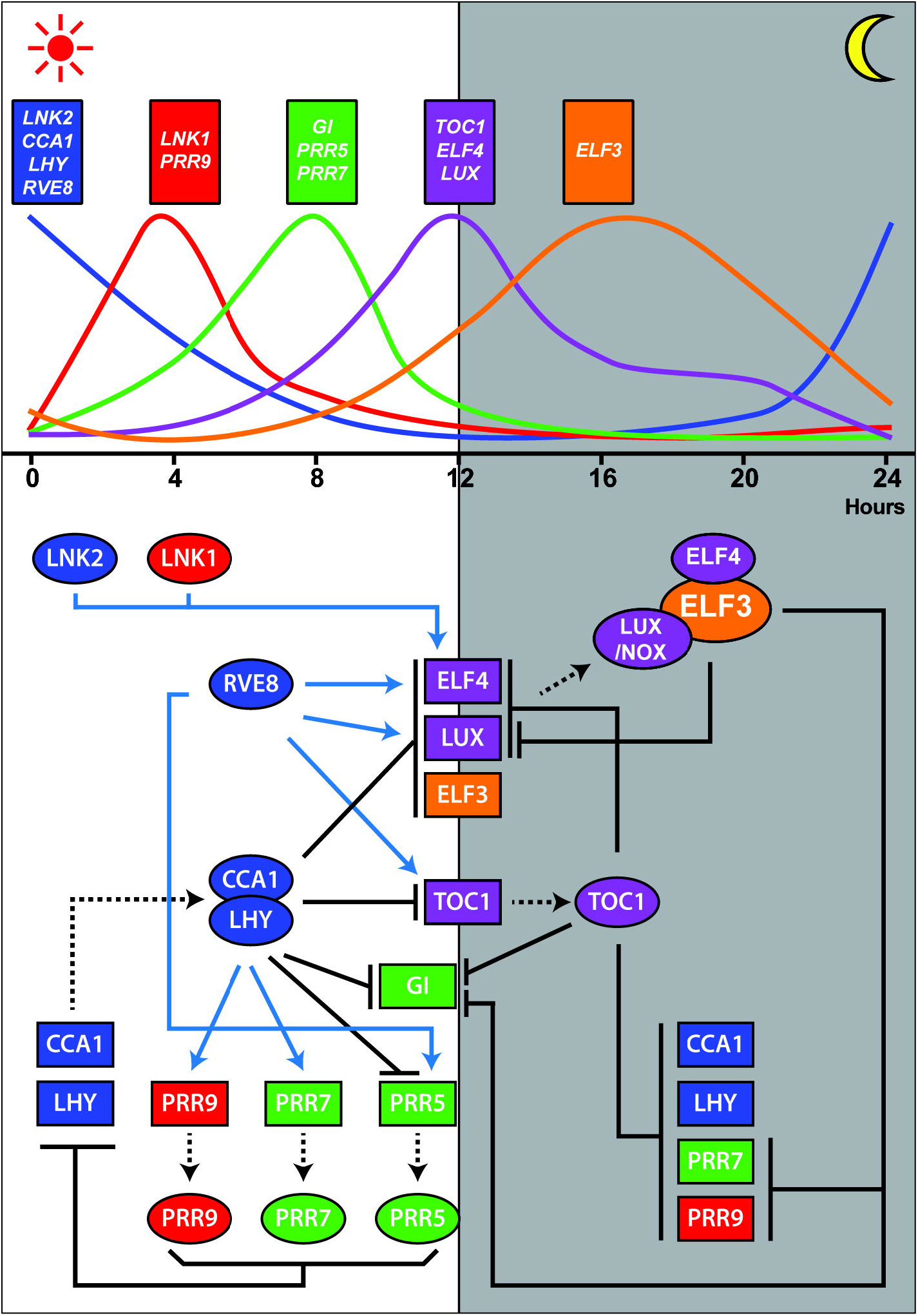
The EC is a key component of the A. thaliana circadian clock. Figure presents the regulatory network of the A. thaliana clock, in which the EC is a critical component of the evening loop, as summarized in the main text and in Box 2. The upper part of the figure illustrates the normalized daily gene expression of key clock genes from publically available time-course microarray data (http://diurnal.mocklerlab.org/) [109]. Genes with similar expression profiles are labeled in the same color. The shade box indicates the night period. The lower part of the figure shows the transcriptional regulation within the *A. thaliana clock*, with blue arrows indicating activation while perpendicular bars indicating repression. Proteins are represented as ellipses and genes as rectangles, with dashed lines indicating translation.

During the last decade, many studies identifying EC-regulated genes and EC-interacting proteins have contributed substantially to our understanding of the circadian network and clock outputs. In this review, we summarize the mechanisms that regulate the accumulation of the EC, and in turn, how the EC modulates the circadian clock, plant growth and development. The role of the EC in connecting different input pathways to the circadian clock and regulation of clock outputs is emphasized. We then focus on reporting recent advances in identifying previously unknown EC-associated proteins, which reveal post-translational connections within the clock pathway and new nodes in clock-controlled outputs. Finally, we provide perspective on utilizing multiple experimental approaches to broaden our knowledge of EC function and the circadian clock in plants.

## Assembly of the EC

The EC is composed of three distinct proteins: ELF4, ELF3, and LUX [12]. *LUX* encodes a GARP transcription factor with a single SHAQYF-type Myb DNA-binding domain [13, 17]. Neither *ELF3* nor *ELF4*, however, encode proteins containing domains with reported function [14-16, 22]. ELF3 directly binds to ELF4 and LUX, functioning as a scaffold within the EC to bring ELF4 and LUX together [12, 23]. Interaction domain mapping using yeast two-hybrid assays indicate that the middle domain of ELF3 (residues 261 to 484) is required for the ELF3-ELF4 interaction [23], while the C-terminus of LUX (residues 144 to 323) is required for interacting with ELF3 [12]. However, the interaction domains of ELF3 for LUX and of ELF4 for ELF3 are not known. ELF4 functions to promote the nuclear localization of ELF3 and presumably the entire EC [23]. LUX directly binds to the LUX binding site (LBS, GAT(A/T)CG) and can recruit the EC to promoters of target genes to suppress their expression [12, 24, 25]. In the absence of LUX, the EC is recruited at lower levels to target promoters through a highly similar GARP transcription factor, NOX (or BOA for BROTHER OF LUX ARRHYTHMO) [12, 25, 26]. NOX expression peaks in the late evening, binds similar DNA sequences, and can directly interact with ELF3 [13, 17, 24, 26]. *NOX*, however, cannot replace *LUX* in sustaining clock activity, as artificial miRNA NOX knock-down lines maintain robust circadian rhythms whereas lux mutants are arrhythmic [24]. Further work is needed to understand how LUX and NOX differ in their activities that contribute to EC function within the clock.

## ELF3, ELF4 and LUX are key cogs in the circadian clock

EC formation appears to follow the daily transcription of its three composite parts, with a single peak tracking dusk under varying day/night conditions [12]. *ELF3, ELF4* and *LUX* cycle under diurnal and free-running conditions (e.g. constant light), suggesting that the clock tightly regulates the oscillation of the EC [12, 13, 15-17, 27]. In turn, the EC regulates the transcription of other key clock genes (discussed below) to maintain proper circadian rhythms [13, 16-18].

### The circadian clock regulates expression of EC components

Supporting the idea that the clock regulates EC transcripts, both *ELF4* and *LUX* promoters contain a cis-regulatory element known as the evening element (EE, AAAATATCT), which is overrepresented in promoters of evening-phased genes [28]. Two morning-phased clock transcription factors CIRCADIAN CLOCK-ASSOCIATED 1 (CCA1) and LATE ELONGATED HYPOCOTYL (LHY) were found to bind the EE and to suppress expression of evening-phased genes [28](Figure 1). Therefore, the nightly peak of ELF4 and LUX expression is likely regulated by CCA1/LHY [17, 29]. Distinct from *ELF4* and *LUX*, the *ELF3* promoter lacks a canonical EE but has one EE-like element (AATATCT) and two CCA1 binding sites (CBS, AA(A/C)AATCT) [30-32]. CCA1 binds to the promoter of ELF3 in the morning to repress its expression, supporting the genetic data showing ELF3 is negatively regulated by CCA1 [33]. Furthermore, a recent systematic approach of identifying genome-wide targets of CCA1 using chromatin immunoprecipitation followed by deep sequencing (ChIP-seq) has shown that CCA1 occupies the promoter regions of all EC components [34, 35].

Other clock factors in addition to CCA1 and LHY contribute to the daily pattern of EC transcript abundance. *ELF4* expression in the afternoon is greatly reduced in the double mutant of two morning-phased genes *LNK1* and *LNK2* (for *NIGHT LIGHT-INDUCIBLE AND CLOCK-REGULATED GENE 1* and *2*), which integrate light inputs into the clock [36]. Additionally, a recent study found that an afternoon-peaking protein REVEILLE 8 (RVE8) antagonizes CCA1 and can activate the expression of *ELF4* and *LUX* through binding to the EE [9, 37]. Near the end of day, an evening-phased central oscillator *TIMING OF CAB EXPRESSION 1* (*TOC1*, also known as *PRR1*) has been found to suppress expression of *LUX* and *ELF4* [32]. Therefore, these data show that various clock components collectively regulate the expression of EC components. However, whether the circadian clock also regulates the protein levels of the EC components through post-translational mechanisms remains unknown.

### Reciprocal regulation of clock genes by the EC

The EC is localized to the nucleus [14, 22, 23, 38], where it functions to mediate nighttime repression of key clock genes *TOC1, LUX, GIGANTEA (GI), PSEUDO-RESPONSE REGULATOR 7* and *9 (PRR7 and 9)* [23, 24, 39-41] and indirectly promotes the expression of the morning oscillators *CCA1* and *LHY* [27, 33, 39, 41] (Figure 1). Two or more of the EC components have been shown by ChIP analysis to associate with the promoters of *PRR7, PRR9, GI* and *LUX* [23-25, 40-42]. LUX also undergoes auto-regulation by binding to its own promoter through the LBS element, suggesting that the EC regulates its own levels through suppressing *LUX* expression [24]. In all cases, EC function is linked to transcriptional repression, as complementation experiments of *lux* mutants have shown that expression of LUX fused to a strong repression domain complements the mutant, while expression of LUX with a strong activation domain does not [12, 24]. The molecular mechanism, however, by which the EC regulates transcription currently remains unclear.

## The EC integrates and transmits multiple light and temperature signals to the clock

Ambient light and temperature are cues that synchronize the internal clock with the external environment in a process called entrainment. It has been shown that both transcript and protein levels of the EC are regulated by light and temperature, and that multiple environmental inputs that communicate timing to the clock are integrated by the EC.

### Light-EC interactions

Since plants are on a constant quest for photons to drive photosynthesis, light signaling influences most processes, including the circadian clock. Both *ELF3* and *ELF4* are regulated by light signaling pathways and induced by light [14, 27]. However, only the molecular mechanism that ties light regulation into *ELF4* expression has been revealed. Three positive transcriptional regulators of the phytochrome A (phyA) light signaling pathway, FAR RED ELONGATED HYPOCOTYL3 (FHY3), FAR-RED IMPAIRED RESPONSE 1 (FAR1) and ELONGATED HYPOCOTYL 5 (HY5) directly bind to the FBS (FHY3/FAR1-binding sites, CACGCGC) and ACE (ACGT-containing elements) cis elements within the *ELF4* promoter to activate its expression during the day [29]. While light regulation of *ELF3* expression in still poorly understood, more is known about post-translational regulation of ELF3 protein levels. For example, the overexpression of the major red light photoreceptor phytochrome B (phyB) stabilizes ELF3 proteins, while a light-regulated E3 ubiquitin ligase CONSTITUTIVE PHOTOMORPHOGENIC 1 (COP1) ubiquitinates ELF3 *in vitro* and negatively regulates the abundance of ELF3 *in vivo* [14, 43, 44]. In addition, a B-box containing transcription regulator that regulates photomorphogenesis, BBX19, physically interacts with COP1 and ELF3 to promote the COP1-dependent degradation of ELF3 [45]. Thus, visible light signaling pathways directly regulate the abundance of the EC through transcriptional and post-translational mechanisms.

No single photoreceptor family in plants has been shown to be necessary for clock entrainment, suggesting a complex interaction between light perception and the oscillator [46]. It has been shown, however, that the EC plays critical roles in regulating light signaling by the clock. ELF3 is a key factor antagonizing light input into the clock, as overexpressing ELF3 attenuates the sensitivity of the clock to both red and blue light-mediated resetting cues [47]. Conversely, a weak allele of *elf3 (elf3-12)* exhibits hypersensitivity to the resetting cue, while a stronger hypomorphic *elf3-7* allele or the null *elf3-1* allele results in severe gating and resetting defects [39, 48]. Similarly, a loss-of-function mutation in *ELF4* also shows a hypersensitivity to resetting and the gating of outputs, suggesting that the entire EC participates in the regulation of light input [49]. The function of the EC as an integrator of light inputs is consistent with protein-protein interactions between the EC and numerous components of the light signaling pathways [14, 38, 43, 44, 50, 51]. The EC also regulates responses to and is regulated by low-intensity non-damaging UV-B light. *ELF4* is highly induced by UV-B light and null mutants of *elf3, elf4* and *lux* mutants exhibit defects in the gating of UV-B responsive gene expression [42, 52]. Consistent with the EC possibly directly regulating UV responses, ChIP analysis showed that ELF4 and LUX are associated with the promoter of a UV-B downstream target gene *EARLY LIGHT INDUCIBLE PROTEIN 1 (ELIP1)* [42]. Although many connections between light signaling pathways and the EC have been observed, how light input from the UV to far red is transmitted into the clock has not yet been solved.

### Temperature-EC interactions

Similar to light, temperature perception is also connected to the EC. Temperature-entrained and dark-grown *elf3* mutant seedlings are arrhythmic, supporting an essential role for ELF3 in integrating temperature cues [53]. Furthermore, ELF3 is required for properly regulating the induction of *GI LUX, PHYTOCHROME INTERACTING FACTOR 4 (PIF4), PRR7* and *PRR9* when Arabidopsis seedlings are shifted to warmer temperatures (22°C to 28°C, or 16°C to 22°C) [40]. The temperature-responsiveness of clock gene expression is abolished in *elf4, elf3,* or *lux* mutants: expression of *GI, LUX, PIF4, PRR7,* and *PRR9,* was found constitutively high regardless of temperature [40]. These findings suggest that the EC integrates temperature input into the clock by regulating the expression of key clock genes (*PRR7/9, LUX* and *GI*) and clock outputs *(PIF4)*. Intriguingly, the association of ELF3 to the promoters of *PRR9, LUX* and *PIF4* is compromised at high temperature, suggesting that temperature might directly regulate EC recruitment to promoters [54].

Cold signals are integrated into the clock through both transcriptional and post-transcriptional mechanisms. A cold-inducible C-repeat (CRT)/drought-responsive element (DRE) binding factor CBF1/DREB1b has been shown to bind the CRT element in the *LUX* promoter and regulate *LUX* expression [55].Decreased intron retention of *ELF3* was also found in the *gemin2-1* mutant, which is known to affect the temperature responsive alternative splicing of several clock genes, including *CCA1*, RVE8, and TOC1 [56]. This observation raises the open question of whether temperature changes directly regulate ELF3 activity through controlling alternative splicing. Similarly, temperature also regulates the alternative splicing of CCA1, which changes the ratio of full-length *CCA1* transcripts in the cellular pool [57]. One could speculate that temperature-regulated alterative splicing of *CCA1* would alter the expression of *ELF4, ELF3,* and *LUX*. Taken together, the transcript and protein abundance of the EC components are regulated at multiple levels by light and temperature pathways.

## EC regulation of clock outputs: growth and flowering

Recent advances have shed light on how the EC connects the clock to output pathways, such as photoperiod-dependent plant growth and flowering. Interestingly, these pathways appear to converge, at least partly, onto the regulation of a small set of transcription factors.

### The EC suppresses expression of PIFs to regulate photoperiodic and thermoresponsive growth

Cellular elongation in the juvenile stem of seedlings, known as the hypocotyl, is highly responsive to changes in both day length and temperature [58, 59]. It is now known that the clock regulates photoperiodic and thermoresponsive growth by tightly regulating key phytochrome-interacting, growth-promoting, bHLH transcription factors known as *PHYTOCHROME INTERACTING FACTORs (PIFs)* [60], particularly *PIF4* and *PIF*5 [12, 40, 61].

Hypocotyl elongation is very sensitive to light conditions and exhibits a non-linear response to the length of night [58]. The circadian clock and light signaling pathways co-regulate hypocotyl elongation to maintain daily growth rhythms in *Arabidopsis*, with the maximal growth rate at dawn in short days [19, 58]. Signals from both the clock and light pathways converge on PIF4 and PIF5 [62, 63]. *PIF4* and *PIF5* protein abundance is post-translationally suppressed during the day mainly by light-activated phyB, while *PIF4* and *PIF5* expression is transcriptionally repressed by the EC during the early evening [12, 19, 58, 64] (Figure 2A). The recruitment of the EC to the promoter region of *PIF4/PIF5* is mediated by LUX and NOX, where NOX acts additively with LUX in regulating plant growth [12]. As the level of the EC decreases as dawn approaches, transcriptional suppression of *PIF4/PIF5* is released [12, 19, 64]. Therefore, the long hypocotyl phenotype of EC mutants can be explained by loss of transcriptional repression at dusk, which leads to the premature accumulation of PIF4/PIF5 proteins and consequently up-regulation of phytohormone-related PIF4/PIF5 target genes, including auxin [65-67]. A recent study also indicates that ELF3 regulates PIF4-mediated plant growth in an EC-independent manner, by directly interacting with PIF4 to prevent the activation of PIF4 transcription targets [44] (Figure 2A). Whether the EC directly regulates the expression or transcriptional activity of other PIFs, such as PIF1, PIF3 and PIF7, remains unclear.

**Figure 2.**
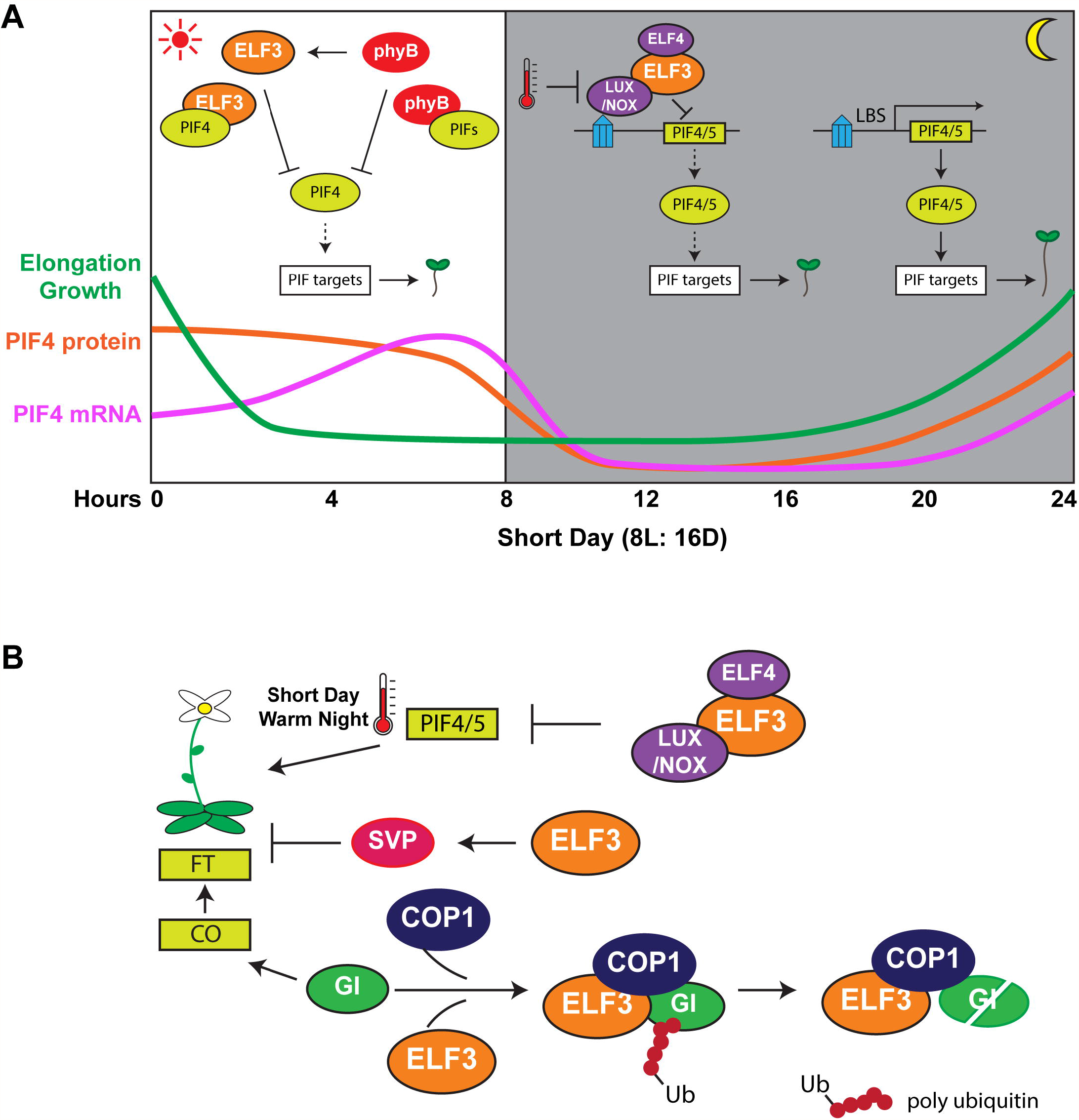
Summary of molecular functions of the EC in regulating growth (A) and flowering (B). (A) The EC suppresses hypocotyl elongation. During the day, ELF3 and phyB suppress PIF4 protein activity and abundance, respectively, while during the night the EC represses expression of PIF4/5. The shade box indicates the night period of the short-day photoperiod condition. LBS=LUX/NOX binding sites. (B) shows the EC-mediated regulation on flowering, including the ELF3-promoted degradation of GI and accumulation of SVP, as well as the EC-mediated suppression on PIF4/PIF5 expression under the short-day, warm night condition. Proteins are represented as ellipses and genes as rectangles.

In addition to shortening days, high-temperature also promotes hypocotyl/petiole elongation and leaf hyponasty [68]. A recent study using 19 *Arabidopsis* natural accessions and quantitative trait locus analysis found that ELF3, but not LUX, contributes significantly to the warm-temperature promoted nighttime elongation growth [54]. A complementary study into natural variation of temperature-responsive growth also provided evidence that a single nucleotide polymorphism (SNP) in the coding region of ELF3 accounts for differences in thermoresponsiveness between two *Arabidopsis* accessions Bay-0 and Sha [61]. Introgression of a single Alanine to Valine substitution at position 362 (converting the Bay-0 ELF3 SNP into the Sha ELF3 SNP) results in an ELF3 hypomorphic allele in the Bay genetic background, which elevates expression of PIF4 and consequently auxin-related PIF4 targets [61, 69]. Therefore, the EC-mediated night-time suppression of *PIF4* expression suggests the EC is essential for both photoperiodic and thermoresponsive growth and connects the clock to both photo- and temperature-responsive outputs.

### The EC regulates flowering

Although two EC components, ELF3 and ELF4, were originally identified in screens for mutants that disrupted flowering timing in *A. thaliana*, how they functioned to regulate the developmental transition to flowering was unclear [16, 70]. One molecular mechanism has been found through studies of ELF3, which functions as a substrate adaptor for the COP1-dependent degradation of GI protein [43, 71] (Figure 2B). Formation of the COP1-ELF3-GI complex destabilizes GI, thereby resulting in reduced expression of flowering-promoting genes *CONSTANS (CO)* and *FLOWERING LOCUS T (FT)* [43]. Additionally, a MADS-box transcription factor SHORT VEGETATIVE PERIOD (SVP), which represses *FT* [72], was shown to directly interact with ELF3 and accumulate in the ELF3 overexpression line [73], consistent with the late-flowering phenotype of ELF3 overexpression lines [43]. It is noteworthy that loss of ELF4 or LUX also causes a similar photoperiod-insensitive early flowering phenotype to *elf3* mutants [16, 17], suggesting that formation of the EC is required to suppress flowering in non-inductive conditions. The EC-target PIF4 also binds to the promoter of *FT* in a temperature-dependent manner and interacts with CO to regulate high-temperature induced flowering under non-inductive short day conditions [74, 75]. Because PIF4 and PIF5 are also required for warm-night induced early flowering [76], it is possible that the EC indirectly regulates flowering by modulating expression of the PIFs or through repressing transcriptional activation through the ELF3-PIF4 interaction.

## EC function in plant species outside of *Arabidopsis*

Orthologs of all the evening complex components, *ELF3, ELF4* and *LUX,* can be found in the reference genomes of land plants lineages from moss *(Physcomitrella patens)* to major crop species [77-87]. Two LUX orthologs ROC15 and ROC75, but not ELF3 or ELF4,have also been found in green algae *Chlamydomonas reinhardtii* [88, 89]. Currently it is unknown if the evening complex forms in any species outside of *A. thaliana*. Analysis of ELF3, however, in *Lemna Gibba* (duckweed) and rice has shown that disrupting ELF3 causes arrhythmia, consistent with its function in Arabidopsis [90, 91]. In addition, recent reports suggest that favorable photoperiodic responses selected through crop improvement are associated with mutations in EC components [78-86]. Using genomic sequencing, the genetic bases of altered photoperiod responses in rice [78-80], pea [81, 82] and barley [83-86] have been uncovered. A number of mutant alleles in *ELF3* and *LUX* orthologs have been identified that result in variation in photoperiod-dependent growth and flowering in these crops. The mutations in *ELF3* or *LUX* are accompanied by changes in circadian rhythms, expression of FT family genes, and GA biosynthesis [78-86]. Subtle alterations in EC function could be used to quickly adapt a crop to a new geographical location by fine-tuning clock, temperature, and light responses. Thus, the EC could be considered as key module for manipulation in order to improve important crop species.

## ELF3 is an important hub of the Arabidopsis protein-protein interaction network

ELF3 is highly interconnected within the circadian network and directly binds to multiple proteins, including phyB, COP1, BBX19, PIF4, ELF4, LUX, NOX, SVP, TOC1, and GI [12, 14, 43-45, 50, 73]**(Table 1)**. This high degree of interconnectivity indicates ELF3 may function as a hub [50] (Figure 3). ELF3 tandem affinity purification coupled with mass spectrometry (AP-MS) was done to identify proteins that directly or indirectly associate with the EC [50]. A curated list of 25 ELF3-associated proteins includes previously reported direct interactors in the circadian clock pathway (ELF4, LUX and NOX) [12] and light signaling pathway (phyB and COP1) [14, 43]. Several new ELF3-associated proteins were also identified, including a family of nuclear kinases (MUT9-LIKE KINASEs, MLK1 to 4), which regulate circadian period and flowering in long days [50, 92]. In addition this study revealed that ELF3 and phyB function as hubs within the EC-phytochrome-COP1 interactome to recruit additional proteins and connect the clock to the light sensory and photomorphogenesis pathways [50]. Among the list, TOC1 was found to be associated with ELF3 *in vivo* and validated as a direct ELF3 interactor in a yeast two hybrid assay [50]. Together with the fact that ELF3 regulates the expression of TOC1 and its target PRR9 [23, 25, 39-41], the TOC1-ELF3 interaction suggests additional mechanisms of EC-mediated regulation within the circadian clock.

**Figure 3.**
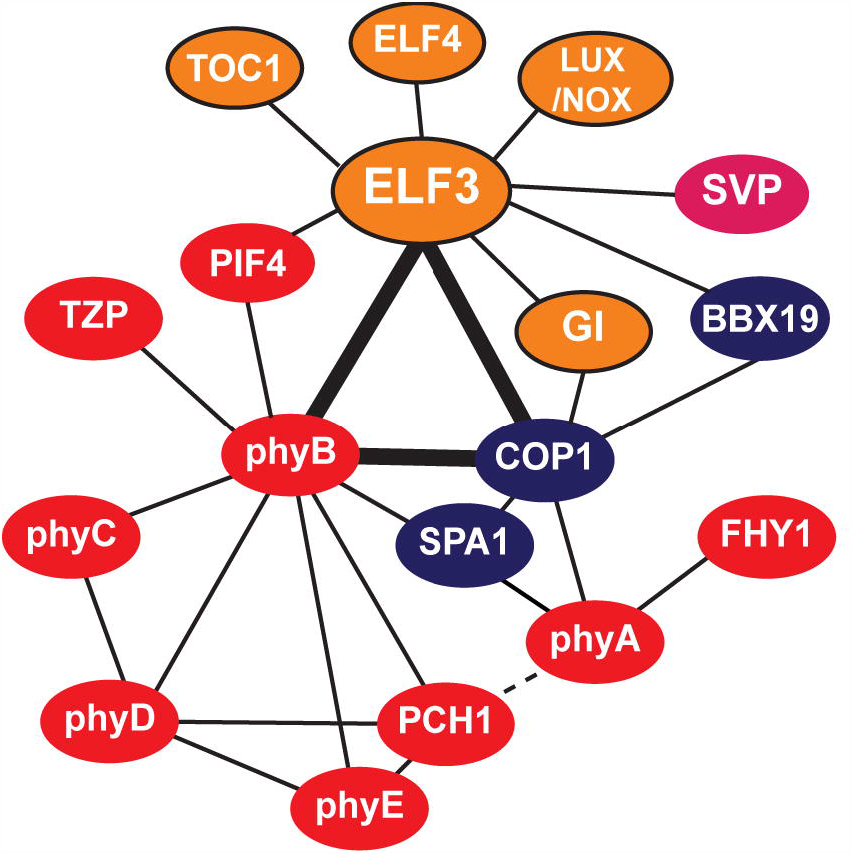
ELF3 is a hub protein of a complicated protein-protein interaction network. Figure shows the structure of the EC-phytochrome-COP1 interactome. Components of the circadian clock pathway are in orange; light signaling components are in red; components of the COP1-mediated photomorphogenesis pathway are in blue. Solid line indicates direct interaction, while dashed line indicates association. The interactions among ELF3, phyB and COP1 are in bold to emphasize that all three proteins are hubs.

**Table 1.** List of proteins directly interacting ELF3

| Protein name | AGI locus | Minimum ELF3 fragment for interaction in Y2H assays | References |
| --- | --- | --- | --- |
| ELF4 | AT2G40080 | AA 261-484 | [12, 23, 50] |
| LUX | AT3G46640 | not tested | [12, 50] |
| NOX | AT5G59570 | not tested | [12, 50] |
| GI | AT1G22770 | AA 1-261 | [43, 50] |
| COP1 | AT2G32950 | AA 1-261 | [43, 50] |
| phyB | AT2G18790 | AA 1-440 | [14, 50] |
| PIF4 | AT2G43010 | AA 442-695 | [44] |
| TOC1 | AT5G61380 | AA 515-695 | [50] |
| SVP | AT2G22540 | not tested | [73] |
| BBX19 | AT4G38960 | not tested | [45] |

It is noteworthy that the powerful combination of genetics and AP-MS can be used to provide insight into the topology of complex protein-protein interaction networks. For instance, the *in-vivo* dependence of phyB for the association of PHOTOPERIODIC CONTROL OF HYPOCOTYL1 (PCH1) and TANDEM ZINC KNUCKLE/PLUS3 (TZP) with the EC was found by comparing AP-MS profiles of EC-associated proteins in WT and *phyB-9* mutant backgrounds [50, 51]. The requirement for phyB suggested that PCH1 and TZP were indirectly recruited to the EC through phyB [50, 51] (Figure 3). Follow up studies determined that TZP and PCH1 directly bind to phyB to regulate either photoperiodic flowering or growth, respectively [51, 93]. Whether their association with phyB and the EC contributes to light input or the clock regulation of outputs is a point of interest for future experiments. Thus, using genetics with AP-MS to probe dependencies in protein complex formation can rapidly lead to testable hypotheses of which proteins are likely functioning together in specific pathways to regulate physiology and development.

## Concluding Remarks and Future Perspectives

Evidence generated from combined genetic, biochemical and molecular approaches has established that the EC functions as an entry point for integrating multiple clock inputs to modulate circadian rhythms and clock output pathways. However, several aspects of EC function and regulation are of great interest for future characterization (see **Outstanding Questions** and below).

First, coupling of tissue-specific circadian clocks have recently been found in plants, suggesting a hierarchical structure of the plant circadian clock [94]. Among several key clock components showing tissue-specific expression patterns, *ELF4* is highly vasculature-enriched (10-fold higher in vasculature) [95]. Additionally, diel cycling of *LUX* expression in vascular tissues peaks at dusk, while in mesophyll and epidermis tissue *LUX* expression shifts to the morning [96]. Currently, all studies of EC function have used whole seedlings. Therefore, whether the composition or activity of the EC is tissue-specific, and how the EC participates in regulating tissue-specific clocks requires further investigation.

Second, recent systematic approaches (ChIP-seq) to identify target genes of several key clock transcription factors (CCA1, TOC1, PRR9, PRR5 and PRR7) [32, 34, 35, 97-99] have expanded our understanding of the complicated regulatory network within the clock and between the clock and its outputs. For example, comparison of TOC1 and PIF3 ChIP-seq datasets found that both proteins are recruited to the promoters of predawn-phased, growth-related genes to simultaneously regulate expression [100]. In addition, TOC1 directly interacts with PIF3 to block transcriptional activation [100], reminiscent of the ELF3-PIF4 interaction [44]. Thus, TOC1-PIF3 interaction may be another point of molecular convergence of the clock and photosensory pathways to control plant growth. Currently, it is an open question if LUX/NOX recruits the EC to PIF4 transcriptional targets to concomitantly repress gene expression similar to the TOC1-PIF3 module. Furthermore, the ELF3-TOC1 interaction may also be important for transcriptional regulation of genes involved in circadian rhythms, growth and flowering. Therefore, genome-wide ChIP-seq analysis aimed at identifying which genes are concomitantly and independently bound by ELF4, ELF3 and LUX/NOX would likely expand our understanding of the role of the EC in regulating gene expression in collaboration with other pathways.

Third, recent proteomic analyses have identified new proteins that associate with ELF3/ELF4 *in vivo*, including an EC-associated clock-output protein (PCH1) and a set of four plant-specific nuclear kinases [50, 51]. The combination of genetics and biochemistry also has shed some light on the hierarchy of the EC-phytochrome-COP1 interactome and the connection between the clock and light signaling pathways [50, 51]. However, the role of most of the newly identified EC connections, both between proteins and pathways, are poorly understood and require further investigation. Does the EC collectively interact with all identified proteins in a “super” complex, in multiple distinct complexes, or as mentioned above, in tissue-specific complexes? Likewise, what are the roles of light and the clock in the regulation of EC-containing protein complex dynamics and function? Furthermore, which proteins or domains participate in regulating transcription by the EC? Clearly, further work is required to fully understand the function of the EC-containing protein networks.

Lastly, selection of altered alleles in EC components may be critical for improving crop production through changing circadian rhythms, flowering responses, and growth, suggesting that the EC is a target for applied crop studies [78-86]. Currently, most of our understanding of EC function comes from experiments done in the model dicot, *Arabidopsis thaliana*. However, it is likely that the EC will interact with different proteins/pathways or regulate new outputs in other species. Therefore, molecular understanding of how EC orthologs function in crop species will be required to elucidate their roles in circadian-, photo-, and thermo-regulation of physiology in crops. Considering that expanding the cultivation areas of crops requires optimization of photoperiod and temperature responses due to climate change, understanding EC function in diverse species could lead to new genetic targets for crop improvement.

## Acknowledgements

We thank Margaret Wilson, Christine Shyu, John Hoyer, Sankalpi Warnasooriya, Katie Greenham, Denise Cunningham, and Josh Gendron for critically reading the manuscript and providing feedback. The Nusinow lab acknowledges support by the National Science Foundation (IOS 1456796). We apologize to labs whose important work was not cited due to space limitations.

